# Super-Resolution Macrophage Imaging via Ultrasound Localization Microscopy and Blinking Nanodroplets

**DOI:** 10.64898/2026.05.07.723418

**Authors:** Saar Gotshal Zahavi, Mike Bismuth, Tiran Bercovici, Tali Ilovitsh

## Abstract

Tracking immune cells deep within living tissue remains a fundamental challenge due to the diffraction-limited resolution of ultrasound imaging and the inability to resolve dense cellular populations. Here, we introduce an intracellular super-resolution ultrasound imaging framework based on stochastic phase-changing nanodroplets (NDs) and ultrasound localization microscopy (ULM). We engineer ∼170 nm perfluorocarbon NDs that undergo reversible, stochastic liquid-gas transitions under acoustic excitation, generating temporally sparse “blinking” signals. Leveraging the intrinsic endocytic activity of macrophages, these NDs are internalized, enabling intracellular contrast generation independent of vascular flow. We validate this approach across imaging scales, from controlled phantoms and *in vitro* systems to *in vivo* tumor models, demonstrating robust intracellular blinking, high cell viability, and consistent super-resolution reconstruction in dense cellular environments. The stochastic blinking of internalized NDs provides the temporal separation required to localize individual sources, overcoming a central limitation of conventional ULM. Following systemic administration, ND-labeled macrophages are tracked *in vivo* after homing to the liver, where super-resolution ULM resolves cellular distributions with a spatial resolution of 26.3 ± 3.2 µm, corresponding to a 6.1-fold improvement over diffraction-limited imaging. This work establishes a previously unexplored paradigm for ultrasound-based intracellular super-resolution imaging, enabling non-invasive visualization of immune cell organization in deep tissue. By introducing spatiotemporally programmable intracellular contrast, this approach expands ultrasound beyond vascular imaging toward functional cellular imaging, with broad implications for immunology, diagnostics, and cell-based therapies.

## Introduction

Macrophages are essential components of the innate immune system, playing a role in tissue homeostasis, immune surveillance, and the regulation of inflammatory responses.^1,2^ Their infiltration into tissues is a key indicator of diverse pathologies, ranging from cancer progression to chronic inflammation.^2–4^ Beyond their diagnostic value, these cells have emerged as key targets and agents for therapeutic interventions, including cancer immunotherapies and targeted drug delivery.^4–6^ Therefore, the ability to track macrophages non-invasively offers vital insights for monitoring the biodistribution and therapeutic efficacy of these treatments.^7,8^ However, this remains a significant challenge. Optical microscopy is limited to superficial tissues, while whole-body methods face inherent trade-offs: magnetic resonance imaging (MRI) is often constrained by low sensitivity, and positron emission tomography (PET) lacks sufficient spatial resolution.^8–10^ Consequently, there is a need for an imaging modality that combines deep tissue penetration with cellular scale resolution.

Ultrasound imaging offers substantial penetration depth for non-invasive body imaging, along with advantages in safety, cost-effectiveness, and accessibility.^11^ However, it is inherently incapable of resolving individual cells due to the diffraction limit.^12,13^ Furthermore, the minimal acoustic impedance mismatch between cells and surrounding tissue results in poor intrinsic image contrast. Therefore, there is a need to develop ultrasound-based methods capable of imaging macrophages *in vivo*. One possible strategy is to exploit the intrinsic endocytic activity of macrophages to internalize ultrasound contrast agents. In ultrasound imaging, microbubbles (MBs) are the most widely used contrast agents.^14^ They consist of a gas core encapsulated by a stabilizing lipid shell and typically have diameters in the range of 1-10 µm. Owing to their strong acoustic scattering, MBs provide sufficient sensitivity to enable detection of single agents *in vivo*. However, due to the diffraction limit, each MB is not resolved as a point object but instead appears as a blurred signal defined by the system’s point spread function (PSF).^15,16^ Previous studies have leveraged MB uptake by macrophages and other endocytic immune cells to enable high-contrast cell tracking using conventional ultrasound.^17–20^ A similar concept has also been explored using smaller contrast agents such as gas vesicles and nanodroplets (NDs).^21–25^ Despite these advances, fundamental spatial resolution constraints remain. Macrophages have a typical diameter of ∼15 µm and often accumulate in dense clusters within tissue.^26^ In contrast, the MB-derived PSF extends over hundreds of microns. As a result, signals from multiple labeled cells spatially overlap, making it infeasible to resolve or isolate individual macrophages based on the ultrasound contrast signal. Ultrasound localization microscopy (ULM) offers a powerful strategy to overcome diffraction-limited resolution, enabling super-resolution imaging by localizing individual MB contrast agents.^16^ In ULM, each MB appears as a diffraction-limited PSF, and its center of mass can be estimated with subwavelength precision. By accumulating localizations over multiple frames, a super-resolved image is reconstructed with spatial resolution improved by an order of magnitude. In conventional implementations, this approach relies on MB motion within the vasculature, where blood flow provides the temporal sparsity and spatial separation required between individual PSFs.^16,27–31^ Extending this concept to cellular imaging, internalization of MBs by macrophages could, in principle, enable their super-resolution visualization. However, this approach faces fundamental limitations. Internalized MBs are confined within cells and do not exhibit sufficient motion to generate the temporal sparsity required for ULM. In addition, macrophages often accumulate in dense clusters, while the acoustic signal from MBs remains continuously “on,” leading to overlapping PSFs from multiple nearby cells.^32,33^ Under these conditions, individual MBs become difficult to reliably isolate or localize, thereby limiting the application of ULM for *in vivo* super-resolution imaging of macrophages.

NDs serve as an alternative class of ultrasound contrast agents, typically composed of a liquid perfluorocarbon (PFC) core stabilized by a lipid shell. Ranging from 100 to 1000 nm in diameter, their liquid state and nanoscale size offer distinct advantages for ultrasound imaging.^27,34^ Upon exposure to an ultrasound pulse, the liquid core of the ND undergoes acoustic droplet vaporization (ADV), transitioning into a highly echogenic gaseous state. The acoustic pressure threshold required for ADV is tunable and depends on parameters such as the ND’s size and the boiling point of the encapsulated PFC.^34–36^ Although utilized for ULM, conventional NDs form relatively stable MBs upon vaporization. Consequently, achieving the spatial sparsity needed for ULM relies on high-intensity, destructive acoustic pulses, a “vaporize-and-destroy” approach that rapidly depletes the contrast agent.^27,37^ To overcome the reliance on destructive pulses or hemodynamics, blinking NDs have emerged as a unique class of contrast agents.^27^ Unlike standard NDs that undergo a permanent liquid-to-gas transition, blinking NDs undergo reversible, stochastic phase changes between liquid and gas states under continuous acoustic exposure or pulse laser irradiation, thereby acting as blinking sources.^38–42^ This stochastic activity provides a mechanism for the temporal separation of overlapping PSFs, allowing ULM methods to localize and reconstruct super-resolution images of dense and stationary agents. However, to date, blinking ND-based ULM has been demonstrated primarily for freely circulating or extracellular agents following systemic administration.^27,38^ The application of this approach to intracellular environments, and in particular for resolving contrast agents internalized within cells, has not been established.

Here, we develop a method for intracellular super-resolution imaging of macrophages *in vivo* using ULM and blinking NDs (Figure 1). By leveraging the natural endocytic activity of macrophages, we labeled these cells with ∼170 nm blinking NDs prior to systemic administration. We show that the stochastic phase-transition dynamics of internalized NDs generate sufficient spatiotemporal sparsity to enable localization and tracking of individual macrophages within dense cellular environments. This approach establishes a previously unexplored paradigm for ultrasound-based intracellular super-resolution imaging, enabling non-invasive visualization of immune cell dynamics deep within living tissues.

**Figure 1:**
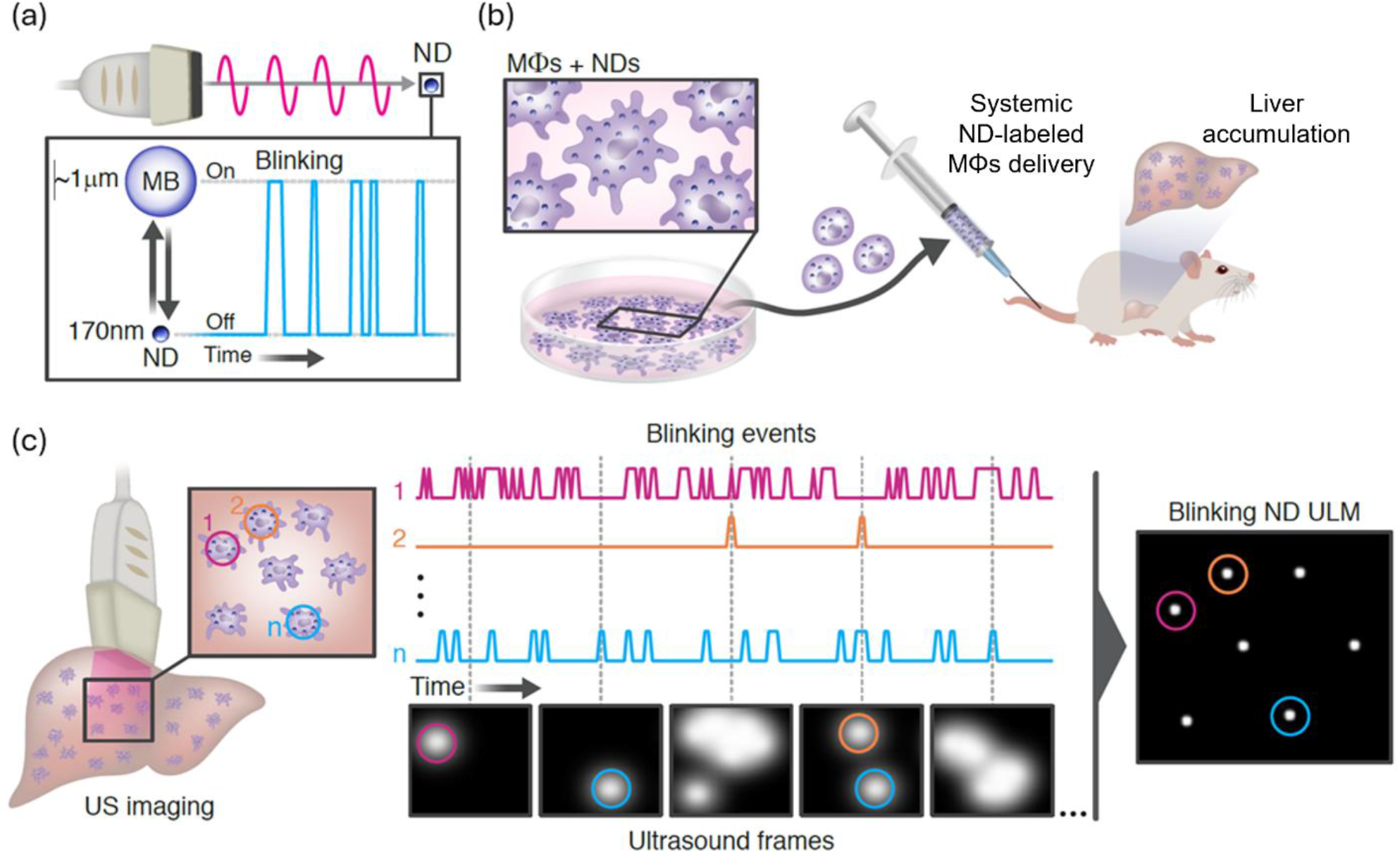
Schematic overview of super-resolution macrophage tracking using blinking NDs. (a) Blinking NDs mode of operation. Under ultrasound exposure, liquid NDs undergo reversable phase transitions into highly echogenic gaseous MBs, generating distinct “on” and “off” temporal signal. (b) Macrophages (MΦs) internalize NDs via endocytosis, the resulting ND-labeled MΦs are systemically administered and subsequently accumulate in the liver. (c) Ultrasound imaging of the liver captures the stochastic blinking events of the ND-labeled MΦs. Fluctuations across sequential frames allow for the spatiotemporal separation of overlapping, diffraction-limited signals. The localizations are then summed to reconstruct a super-resolution ULM map of the macrophages.

## Results and discussion

### ND characterization and acoustic blinking activation

NDs were synthesized using a microfluidic Herringbone Mixer (Figure 2a) to precisely control particle size and concentration, yielding an average diameter of 170 ± 50 nm (Figure 2b). To evaluate the effect of shell composition on signal stability, two formulations were prepared, with and without the surfactant Pluronic (L61). Transmission electron microscopy (TEM) confirmed that both ND formulations shared a similar spherical morphology (Figure 2c, d). To evaluate their acoustic response, NDs were examined *in silico* within agarose gel phantoms. Upon exposure to ultrasonic waves, the NDs underwent stochastic blinking activation. Temporal analysis of this activity revealed unique patterns for each droplet (Figure 2g), with an average frequency of 2.09 ± 1.93 Hz for NDs synthesized with Pluronic (Figure 2e). These temporal signatures, combined with the spatial peaks identified from the maximum intensity projection (MIP) of the subtracted frames (Figure 2f), enabled the accurate detection of distinct NDs.

**Figure 2:**
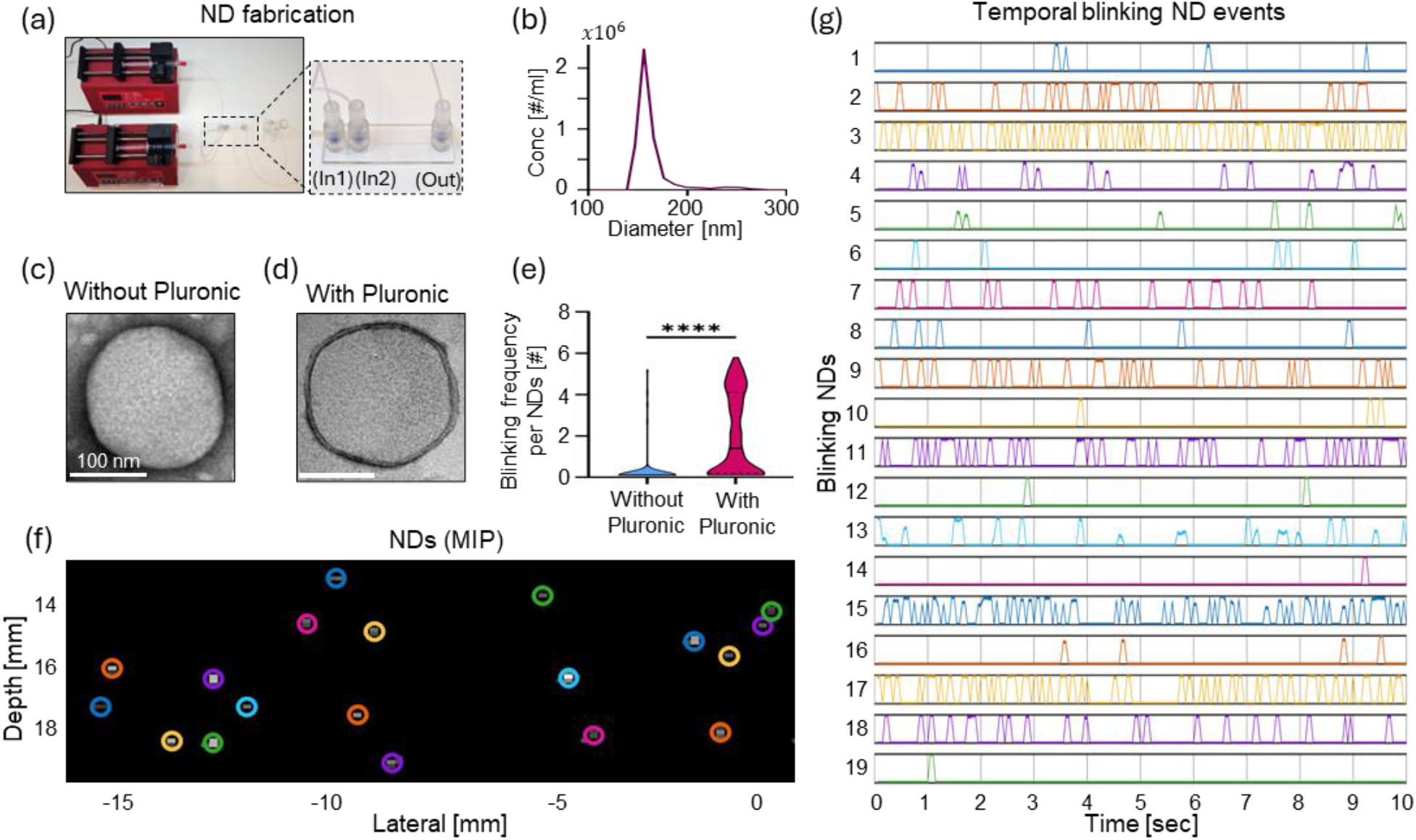
ND fabrication, size distribution and acoustic blinking characterization. (a) ND fabrication via microfluidic chip. (b) Typical size distribution of NDs with Pluronic (L61). (c, d) TEM of NDs. (c) Without Pluronic, and (d) with Pluronic. (e) Blinking frequencies for NDs formulated with and without Pluronic (L61). Unpaired t-test, *n* = 8. (f) MIP for NDs with Pluronic *in silico*, marked by colors. (g) Individual ND blinking patterns for corresponding NDs as marked in (f). NDs are marked from left to right (f) correspond to (g) from top to bottom. Adjusted p value was \*\*\*\**p* < 0.0001. All data are plotted as the median ± quartiles.

However, a significant difference in signal stability was observed between the two formulations. While NDs lacking Pluronic failed to demonstrate sustained activity, the Pluronic-modified NDs exhibited robust, repetitive blinking, with individual droplets undergoing up to 60 blinking cycles during the 10-second recording period (Figure 2e). This absence of repetition in the non-Pluronic group is attributed to lower re-condensation stability, a finding consistent with previous studies demonstrating that Pluronic incorporation enhances ND blinking probability.^38,43^ Consequently, the Pluronic-modified formulation was selected for all subsequent experiments.

### MΦ internalization of NDs for cellular labeling

To internalize the NDs, we leveraged the natural endocytic activity of macrophages. NDs were introduced to the cell culture media, and intracellular uptake was evaluated following incubation periods of two and four hours (Figure 3a, b). Microscopy analysis demonstrated significant ND accumulation within the cells at both time points. An average of 5.37 ± 6.7 NDs per cell was observed after a two-hour incubation period, and 7.6 ± 6.47 after four hours. As no significant difference in uptake efficiency was observed between the two durations, a two-hour incubation period was selected for all subsequent experiments. Furthermore, fluorescence imaging of the labeled cells embedded in an agarose gel phantom revealed a uniform spatial distribution (Figure 3c). These results confirm the successful formation of ND-labeled macrophages, establishing the basis for investigating their acoustic blinking properties.

**Figure 3:**
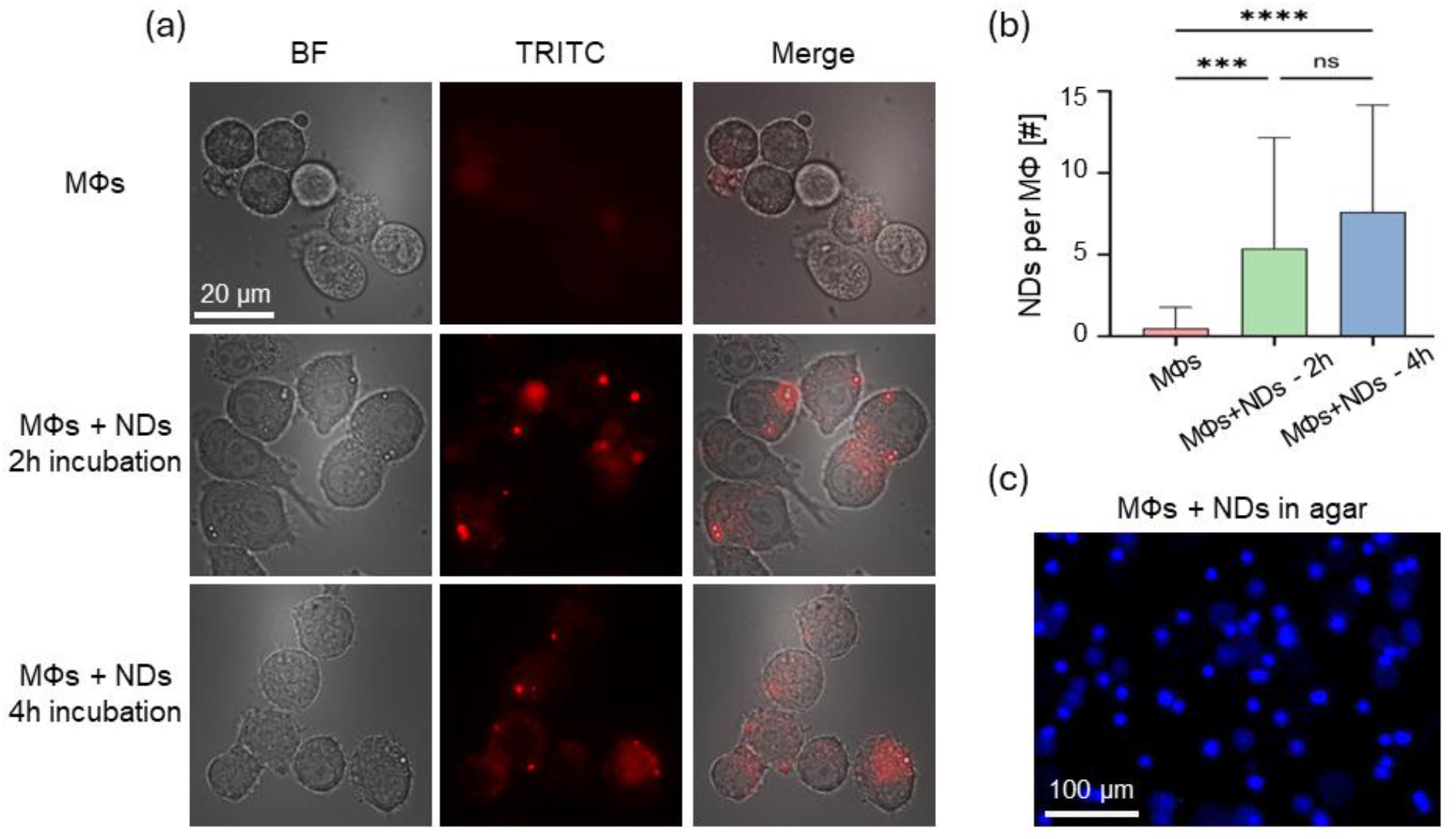
MΦs load blinking NDs, forming ND-labeled MΦs. (a) Fluorescence microscopy images comparing untreated MΦs with MΦs incubated with fluorescent red NDs (DiI) for two and four hours (60x magnification). (b) Number of NDs internalized in MΦs, per cell cross-section. One-way ANOVA with Tukey’s multiple comparison test. (c) Fluorescence microscopy of ND-labeled MΦs in agarose gel, stained blue with DAPI (4x magnification). Adjusted p values were \*\*\**p* < 0.001, and \*\*\*\**p* < 0.0001. *n* = 4. All data are plotted as the mean ± SD.

### Ultrasound induced intracellular ND blinking in MΦs

Prior to cellular imaging, we assessed the impact of the biological labelling conditions on ND acoustic performance. Comparisons between fresh NDs and those subjected to a two-hour incubation at 37°C revealed no significant degradation in activity. The number of detected blinking NDs was comparable between the two groups (Figure 4a), indicating that the incubation period required for macrophage uptake does not compromise the NDs’ acoustic sensitivity. For both conditions, the blinking initiated at a peak negative pressure (PNP) of ∼2 MPa, with the number of events increasing until reaching a maximum at 5.2 MPa, the upper pressure limit of the L12-3v transducer. Importantly, viability assays conducted following ultrasound exposure confirmed that ND-labeled macrophages maintain high cell viability of 94.43% ± 8.83% (Figure 4b), demonstrating that the intracellular blinking activations do not induce cytotoxicity.

**Figure 4:**
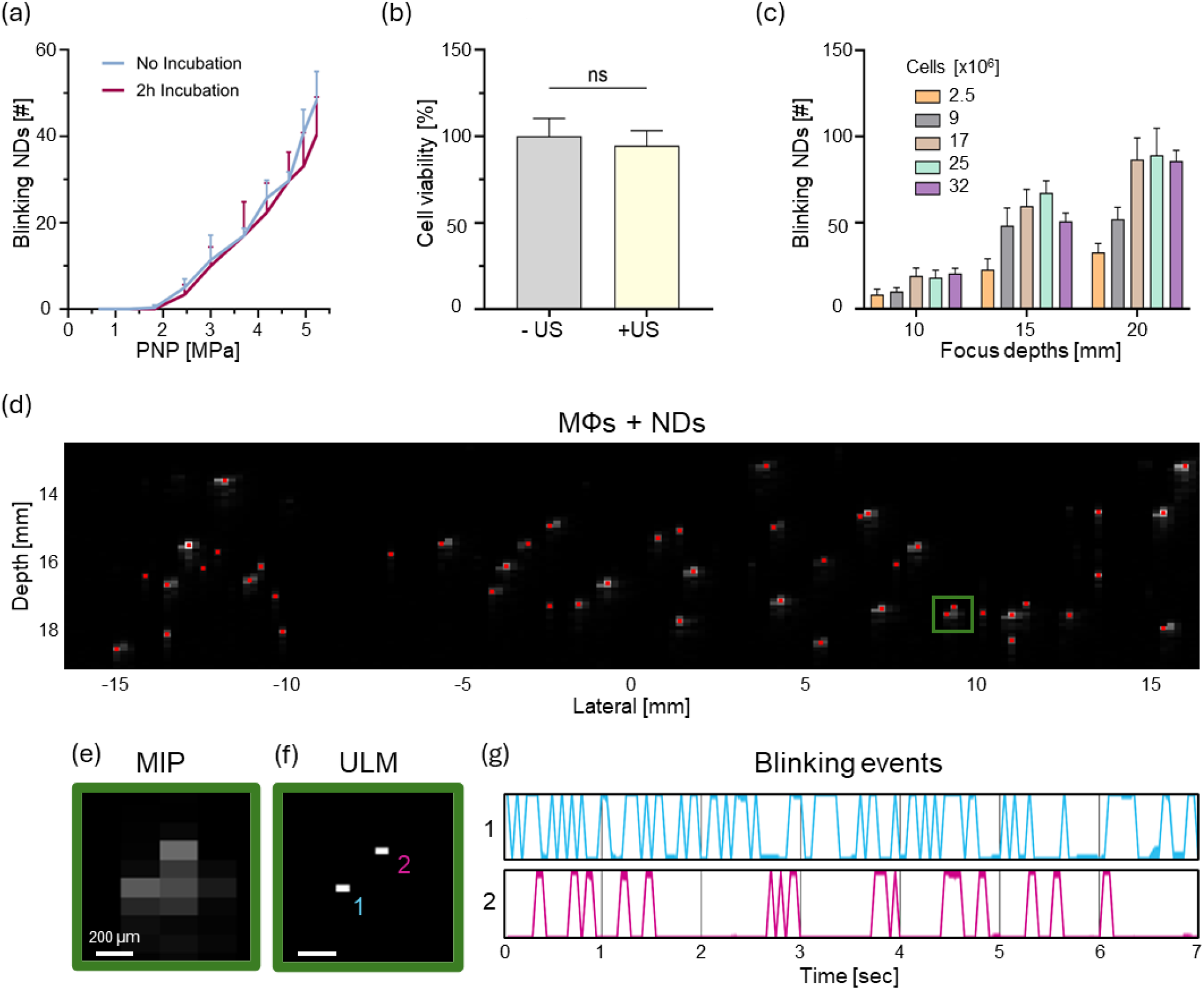
Acoustic activation of ND-labeled MΦs *in vitro*. (a) Acoustic pressure and incubation effects detection of NDs, *n* = 3. (b) Viability of ND-labeled MΦs after US activation. Unpaired t-test, *n* = 6, p>0.05. (c) Detected ND-labeled MΦs for different cell amounts, *n* = 6. (d) MIP for ND-labeled MΦs *in vitro*, ULM localizations marked in red. (e, f) Separation of ND-labeled MΦs, MIP (e) and ULM image (f) of the area marked green in (d). (g) Corresponding blinking events for the ND-labeled MΦs in (f). All data are plotted as the mean ± SD.

To verify that the detected blinking signals originated specifically from internalized NDs, we assessed the blinking signal across phantoms containing increasing concentrations of ND-labeled macrophages. Although a constant initial concentration of NDs was applied to all groups, the number of detected blinking events increased proportionally with the number of labeled cells (Figure 4c). This trend, observed consistently across three different focal depths, strongly supports the hypothesis that the blinking signals arise from endocytosed NDs internalized within the cells, rather than from non-specific background particles.

Next, the stochastic nature of these blinking signals was leveraged to achieve super-resolution ULM (Figure 4d). To demonstrate this, we analyzed two adjacent ND-labeled macrophages that appeared as a single, unresolvable cluster in the standard MIP image (Figure 4e). By analyzing their temporal fluctuations (Figure 4g), we successfully distinguished the neighboring sources. Because the two NDs exhibited distinct blinking patterns, with blinking frequency of 4.86 Hz and 2.14 Hz, respectively, they could be separated and localized in different frames, achieving sub-pixel resolution (Figure 4f). Applying this ULM algorithm across the entire field of view enhanced the spatial resolution by a factor of 2.4 over the standard MIP images, enabling the visualization of adjacent macrophages at a resolution of 182.7 ± 10.8 µm. Importantly, this improvement reflects the specific spatial distribution of cells within the phantom, where macrophages were relatively sparsely packed. As such, the measured factor should not be interpreted as an intrinsic limit of the method. In more densely populated environments, such as *in vivo* tissues where macrophages are expected to be in closer proximity, the increased spatial overlap combined with stochastic blinking is expected to further enhance the achievable resolution. These results highlight that the stochastic intracellular blinking provides the temporal sparsity required to resolve dense, overlapping targets, establishing a foundation for super-resolution imaging of cellular populations *in vivo*.

### *In vivo* ULM of breast cancer tumors using ND-labeled MΦs

We next sought to extend this approach to *in vivo* imaging in a murine model of breast cancer. Given the superficial nature of breast tumors, we aimed to exploit higher ultrasound frequencies to improve spatial resolution. To this end, we transitioned to the L22-14vX transducer (center frequency 18 MHz). As a first step, we verified that stochastic blinking of NDs could be efficiently detected at this higher frequency. An average of 54 ± 8.3 blinking NDs were detected using the L22-14vX, representing a 1.97-fold increase compared to the ones detected with the L12-3v (Figure 5a). Notably, this robust activation was achieved at a PNP of only 1.2 MPa, corresponding to a mechanical index (MI) of 0.3. This demonstrates that significant blinking activation is not dependent on high acoustic pressures and can be induced efficiently at clinically safe, low-MI levels.

**Figure 5:**
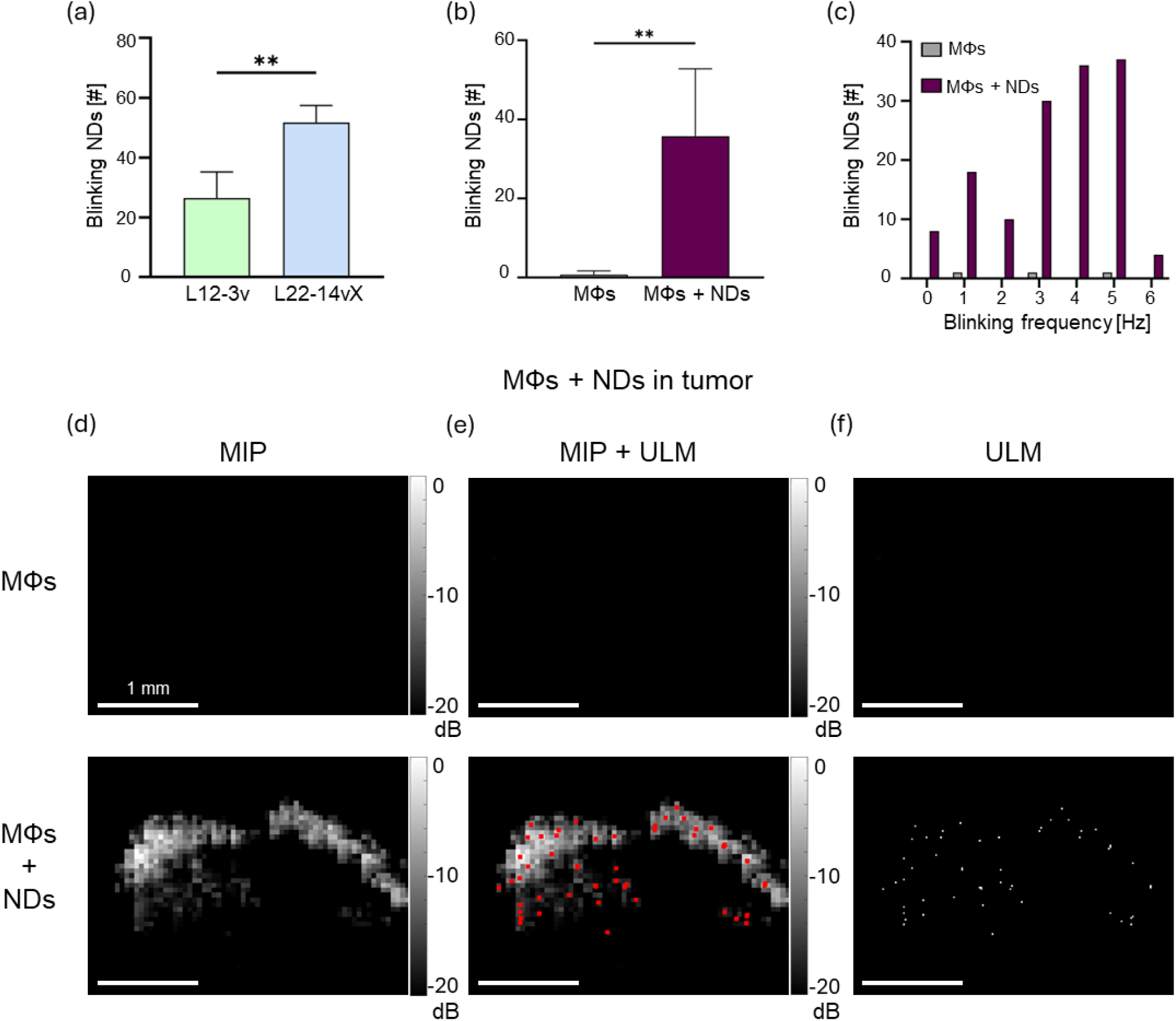
ULM of ND-labeled MΦs in a breast cancer mouse model. (a) Number of blinking NDs detected with the transducers L12-3v (5.1 MPa) and L22-14vX (1.2 MPa). Unpaired t-test, *n* = 7. (b) Detected ND-labeled MΦs *in vivo*. Unpaired t-test, *n* = 4 tumors. (c) Blinking frequencies for ND-labeled MΦs *in vivo, n* = 4 tumors. (d, e, f) MIP and ULM images of MΦs in the tumor (top) and ND-labeled MΦs (bottom); (d) MIP. (e) MIP with ULM localizations labeled in red. (f) ULM. Adjusted p values were \*\**p* < 0.01. All data are plotted as the mean ± SD.

Following this validation, we evaluated the feasibility of tracking immune cells in a localized tumor model, prior to investigating systemic distribution. ND-labeled macrophages were administered into the tumor, followed by ultrasound imaging. Imaging of a control group injected with unlabeled macrophages was also performed for comparison. We detected an average of 35.75 ± 17.06 blinking events from the ND-labeled macrophages *in vivo*, compared to 0.75 ± 0.96 in the unlabeled control group (p < 0.01, Figure 5b). The minimal background signal in the control group is likely attributable to minor physiological motion artifacts, such as respiration or cardiac pulsation. Furthermore, we observed blinking frequencies ranging from 0 to 6 Hz (Figure 5c), confirming the stochastic nature of the phase-change events necessary to achieve the temporal separation required for ULM. In the MIP images (Figure 5d), the dense accumulation of cells appears as a blurred continuous signal. By localizing the individual blinking events (Figure 5e), the reconstruction of the stochastic signals provided a super-resolved image with a spatial resolution of 30 ± 5.6 µm *in vivo* (Figure 5f), representing a 4.5-fold enhancement compared to the diffraction-limited MIP images. These results validate that the ND-labeled macrophages can be robustly activated and resolved through biological tissue, successfully translating this tracking framework to an *in vivo* setting.

### ULM imaging of ND-labeled MΦs in the liver

Following *in vivo* tumor imaging, we next sought to validate systemic administration and assess the biodistribution of ND-labeled macrophages in a highly perfused organ. To this end, we evaluated their accumulation in the liver. ND-labeled macrophages were administered intravenously and allowed to circulate for four hours to facilitate biodistribution and active liver homing *in vivo*. Due to its proximity to the lungs and heart, the liver was imaged ex vivo to minimize respiratory and cardiac motion artifacts, ensuring that detected blinking signals originate from NDs rather than tissue motion. The brain was also assessed as a negative control organ, given that macrophages are not expected to cross the intact blood-brain barrier (BBB).

Ultrasound imaging of the liver revealed robust blinking activity from the internalized NDs. These temporal events remained stable, exhibiting up to 60 blinking cycles during the 10-second recording period (Figure 6e), a performance comparable to the *in vitro* observations (Figure 2e). This confirms that the ND-labeled macrophages maintained their acoustic activity throughout the injection, circulation, and tissue accumulation processes. Leveraging these signals, ULM resolved diffraction-limited MIP features into super-resolved maps (Figure 6a-c), achieving a spatial resolution of 26.3 ± 3.2 µm. This represents a 6.1-fold improvement over MIP. Blinking activity was minimal in the brain (Figure 6d).

**Figure 6:**
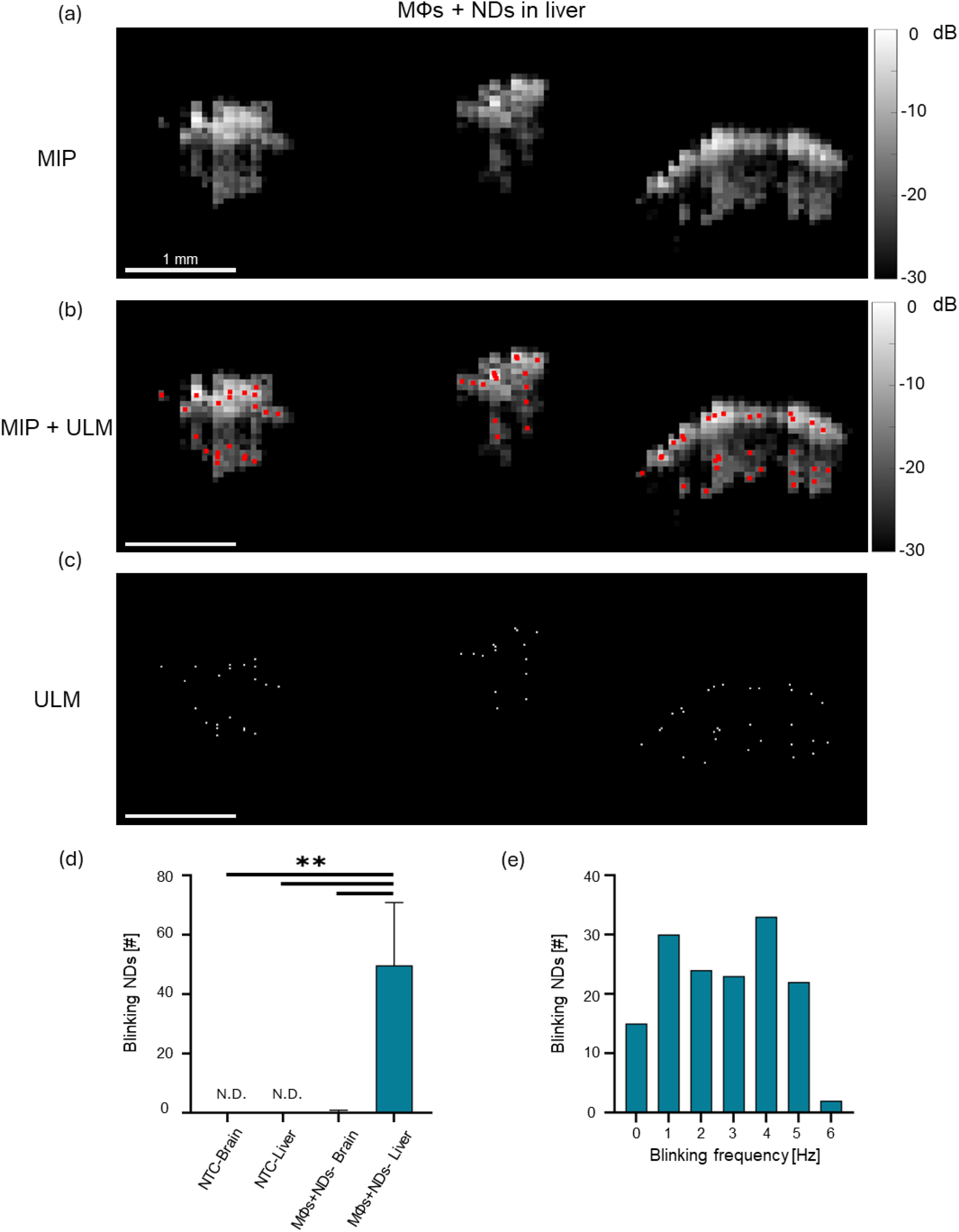
ULM of ND-labeled MΦs in the liver. (a) MIP. (b) MIP with ULM localizations labeled in red. (c) ULM. (d) Detected ND-labeled MΦs in the liver. One-way ANOVA with Tukey’s multiple comparison test. (e) Blinking frequencies for ND-labeled MΦs in the liver. Adjusted p values were \*\**p* < 0.01, *n* = 3. All data are plotted as the mean ± SD.

The ULM images revealed that macrophages accumulate within the liver in distinct clusters of approximately 1-2 mm, with three such clusters observed within a single image plane (Figure 6a-c). This localized accumulation pattern exceeds the dimensions of individual murine lobules (∼0.5 mm), suggesting that the macrophages distributed across supralobular vascular domains spanning multiple adjacent functional units. These domains comprise sinusoidal networks where resident macrophages naturally accumulate to filter the blood.^44,45^ This anatomical correspondence demonstrates that the method reflects the functional distribution of macrophages within the organ, marking a significant step towards super-resolution intracellular immune cell tracking via ultrasound imaging.

While this study establishes a robust framework for *in vivo* tracking of macrophages, several limitations remain. Deep-tissue imaging in dynamic organs is susceptible to motion artifacts; therefore, systemic biodistribution was assessed *ex vivo* to ensure reliable localization. Future studies will incorporate motion compensation algorithms to enable fully in vivo implementation of the method. In addition, the performance of the framework at higher cellular densities remains to be systematically explored. The observed 6.1-fold resolution improvement reflects the specific *in vivo* cell distribution and may vary in denser populations, although such conditions are not directly controllable *in vivo*. Furthermore, the stochastic blinking behavior of NDs is strongly influenced by US excitation parameters and imaging frame rate, and its tunability remains to be optimized. Finally, variability in intracellular ND uptake may introduce heterogeneity in signal intensity across cells.

Future work will focus on motion correction strategies, improved control of blinking dynamics, and integration with volumetric US imaging to enable fully *in vivo* super-resolution mapping of immune cell dynamics.

## Conclusions

This study establishes an intracellular super-resolution US imaging framework that overcomes the diffraction limit to enable non-invasive tracking of macrophages in deep tissue. By leveraging blinking NDs as intracellular contrast agents, we demonstrate ULM without reliance on vascular flow, addressing a central limitation of conventional approaches. The stochastic blinking behavior of NDs was preserved following cellular internalization and across imaging configurations using transducers operating at distinct center frequencies, confirming the robustness and translatability of the platform. We validated this framework across imaging scales, from *in vitro* systems to *in vivo* tumor models, where dense macrophage accumulations were resolved with enhanced spatial resolution. Following systemic administration, ND-labeled macrophages were successfully tracked after homing to the liver, achieving a 6.1-fold improvement in spatial resolution and reaching 26.3 ± 3.2 µm. Together, these results demonstrate that intracellular blinking provides the temporal sparsity required for super-resolution imaging in dense cellular environments, establishing a new paradigm for US-based cellular imaging. This capability advances US from vascular imaging toward functional, cell-resolved imaging, with potential applications in non-invasive diagnostics and the monitoring of cell-based therapies.

## Methods

### ND preparation and characterization

NDs were synthesized via a microfluidic fabrication process by mixing organic and aqueous solutions within distinct microchannels under controlled laminar flow. The organic solution comprised the lipid powders distearoylphosphatidylcholine (DSPC) (Sigma-Aldrich) and 1,2-distearoyl-sn-glycero-3-phosphoethanolamine-N [methoxy(polyethylene glycol)-2000] (ammonium salt) (DSPE-PEG2K) (Sigma-Aldrich). The lipid powders were dissolved in ethanol at a molar ratio of 90:10. For Pluronic-modified NDs, poly(ethylene glycol)-block-poly(propylene glycol)-block-poly(ethylene glycol) (Pluronic L61) was added to the solution to achieve a molar ratio of 90:10:0.1 (DSPC:DSPE-PEG2K:Pluronic L61). In both cases, the final DSPC concentration was 1 mg/mL. The solutions were heated to 40°C and sonicated for five minutes. Subsequently, 1,1′-Dioctadecyl-3,3,3′,3′-tetramethylindocarbocyanine perchlorate (DiI) (Sigma-Aldrich) was dissolved in the organic solution, and the perfluorocarbon perfluoro-n-hexane (perfluorohexane) (SynQuest Laboratories) was added to a final concentration of 2% (v/v) followed by vortexing to ensure homogeneity. The aqueous solution was made using water, propylene glycol (PG, Sigma Aldrich, Milwaukee, WI) and glycerol (Gly, Acros Organics) at a volume ratio of 7:2:1, as described previously.^38^

The ND structure was formed using a microfluidic chip (0.2 mm channel width, Herringbone Mixer Glass Chip, Darwin Microfluidics, UK) and two syringe pumps (NE-300 InfusionONE Syringe Pump, Darwin Microfluidics, UK). The aqueous solution was pumped into the chip at inlet 1 at a velocity of 2 mL/min, while the organic solution was introduced simultaneously through inlet 2 at a velocity of 500 µl/min (Figure 2a).

The ND suspension was dialyzed against phosphate-buffered saline (PBS) using a cellulose membrane (D9777, Sigma-Aldrich) to remove unencapsulated components and ethanol. The dialysis was performed in two consecutive cycles of 1.5 hours each, maintaining a 1:20 (v/v) sample-to-buffer ratio. The NDs were stored at 4 ºC.

The size distribution and concentration of the NDs were measured using a particle counter system (AccuSizer FX-Nano, Particle Sizing Systems, Entegris, MA, USA). The morphology of the NDs was further examined using TEM (JEM-1400Plus, JEOL, Tokyo, Japan) as previously described.^46^

### Ultrasound acquisition

Ultrasound imaging and ND activation were performed using a programmable ultrasound system (Vantage 256, Verasonics Inc., Redmond, WA, USA) equipped with L12-3v and L22-14vX linear array transducers (Verasonics). Ray line sequences of focused ultrasound pulses were implemented in MATLAB (R2025a, MathWorks, Natick, MA). The number of active elements was calculated automatically based on the focus depth and F-number (Table 1). In each experiment, an agarose phantom was positioned such that the region of interest was aligned with the focal spot of the transducer. Center frequencies of 8 MHz (L12-3v) and 18 MHz (L22-14vX) were utilized for both the application of acoustic pressures to activate the blinking NDs and for acquiring B-mode ultrasound images. Data was recorded in blocks of 200 continuous frames at a frame rate of 20 Hz. The PNPs of both transducers were calibrated with a needle hydrophone (NH0200, Precision Acoustics, Dorchester, UK) in a degassed water tank.

**Table 1:** Ultrasound parameters and specifications.

| Transducer | Frequency [MHz] | Pulse length [cycle] | F-number | Focus depth [mm] | Active elements [#] | PNP [MPa] | MI |
| --- | --- | --- | --- | --- | --- | --- | --- |
| L12-3v | 8 | 1 | 2 | 10-20 | 25-50 | 3.9-5.2 | 1.4-1.7 |
| L22-14vX | 18 | 1 | 0.5-0.8 | 6-10 | 100-128 | 0.6-1.2 | 0.1-0.3 |

### ULM image processing

Data analysis was conducted on the recorded 200-frame B-mode sequences. A differential frame-by-frame subtraction process was used to distinguish blinking ND signals from the surrounding static tissues. Data was normalized based on the global minimum and maximum intensities of each experiment. The MIP of the blinking events was calculated to provide a summarized image containing all NDs that appeared throughout the recording.

Blinking NDs were detected via spatial and temporal analysis. First, peak detection was conducted using the “Imregionalmax” function to detect local maxima on the MIP image. Second, each spatial peak was analyzed temporally, and the blinking events were detected with the “findpeaks” function. Two spatial peaks within a 10-pixel distance that shared a temporal similarity greater than 75% were considered duplicates; in such cases, the ND with the lower blinking intensity was removed.

Localization of NDs was then performed for every frame separately, within a 6x6 pixel window centered around each local maximum using an elliptical Gaussian fitting model. The confidence of each localization was calculated as the R^2^ value. For each ND, the localization with the highest confidence was chosen, subject to a confidence threshold of 0.65. The final ULM images were reconstructed by summing all valid localization points detected in the sequence. All data processing was conducted in MATLAB.

Furthermore, to estimate the resolution enhancement, we analyzed dense clusters where individual signals were resolvable. The mean distance between the closest distinguishable peaks and standard deviation were calculated in ImageJ for both the MIP and ULM images. The ratio between the mean values (MIP/ULM) defined the fold-improvement in spatial resolution. All resolution enhancement calculations were analyzed in triplicates.

### Ultrasound imaging of blinking NDs *in vitro*

To evaluate acoustic performance, both ND formulations were dispersed separately into liquid 1.5% (w/v) agarose solutions at 55°C to reach final concentrations of 8.8×10^8^ NDs/mL. The solutions were then poured into custom molds and cooled to room temperature for gel formation. The acoustic blinking ND activation and data acquisition were achieved using the L12-3v transducer. For all subsequent experiments, the Pluronic-modified formulation was utilized.

The effect of thermal incubation and the acoustic pressure was assessed by comparing NDs maintained at 4°C with those incubated at 37°C for two hours prior to gel embedding, with both groups yielding final concentrations of 8.8×10^8^ NDs/mL. Acoustic blinking ND activation and data acquisition were performed using the L12-3v transducer at decreasing PNPs ranging from 0-5.2 MPa.

### Macrophage cell culture

RAW 264.7 murine macrophages were used as model cells for intracellular ultrasound imaging. The macrophages were cultured in high-glucose Dulbecco’s Modified Eagle Medium (DMEM) supplemented with 10% (v/v) fetal bovine serum (FBS), 1% (v/v) penicillin-streptomycin, and 1% (v/v) L-glutamine. Cells were maintained at 37°C in a humidified incubator with 5% CO_2_ until they reached approximately 80% confluency on the day of the experiment.

### Preparation of ND-labeled MΦs and validation

Macrophages were grown in 6-well plates or T75 tissue culture flasks and maintained as described above. On the day of the experiment, the culture medium was replaced with a suspension of NDs in DMEM at a final surface concentration of 8×10^8^ NDs/cm^2^ of tissue plate’s surface. For endocytic uptake of NDs, the cells were incubated with the ND suspension for either two or four hours for the validation experiment, and two hours for all subsequent procedures. Following incubation period, the ND suspension was removed, and the cells were rinsed to remove non-internalized NDs. For experiments using 6-well plates, the cells were rinsed five times with 1 mL DMEM, and for experiments using T75 flasks the rinse included five washes with 5 mL DMEM and two additional washes with 25 mL PBS.

To validate intracellular uptake, cells were imaged using an Echo Revolution microscope (ECHO, San Diego, CA, USA) equipped with a 60x oil immersion lens. To accommodate the short working distance of the immersion lens, ND-labeled macrophages were dissociated with a cell scraper and transferred to glass coverslips. The cells were incubated for an additional hour to ensure adherence prior to imaging. Intracellular NDs were identified via fluorescence imaging of the encapsulated DiI dye using the TRITC channel, overlaid with brightfield (BF) images. The number of NDs within each cell was quantified manually from the resulting fluorescence images.

### ND-labeled MΦs’ viability after US activation

For viability assays, ND-labeled macrophages were prepared as described above in 6-well plates along with a control group of macrophages without NDs. The cells were then collected via dissociation with TrypLE Express (Gibco Corp,12604-013, Grand Island, NY, USA) and resuspended in 500 µL of degassed DMEM.

Ultrasound acquisition to induce ND blinking activation was applied to the cell suspensions within a custom 1.5% (w/v) agarose phantom. The phantom was cast in a mold (65 mm × 25 mm × 20 mm) containing a centrally positioned aluminum rod (15 mm height, 6 mm diameter) to create a sample well. After removal from the mold, the phantom was aligned with the focal zone of the L12-3v transducer. The rod-shaped cavity was filled with the cell suspension, which was gently pipetted to maintain cell distribution during the one-minute ultrasound acquisition at 5.2 MPa. Following exposure, the cell solutions were re-seeded in 6-well plates and maintained at 37°C for 24 hours to monitor post-activation recovery. Finally, the cells were dissociated with TrypLE and quantified using a CellDrop automated cell counter (CellDrop, DeNovix Inc., Wilmington, DE, USA).

### Ultrasound imaging of ND-labeled MΦs *in vitro*

To confirm that the acoustic blinking signals originated from within the macrophages, ultrasound imaging was conducted on five agarose gels containing varying cell concentrations. Macrophages were seeded in five T75 flasks at varying initial densities to reach a range of confluences (40%-100%) by the day of the experiment. All flasks were treated with a constant ND surface concentration of 8×10^8^ NDs/cm^2^. The cells were dissociated with a cell scraper in 7 mL DMEM, quantified, centrifuged and resuspended in 1 mL of degassed PBS. The resulting ND-labeled macrophage suspensions (containing 2.5, 9, 17, 25 and 32×10^6^ cells) were subsequently dispersed separately into 20 mL liquid 1.5% (w/v) agarose solutions at 55 °C, poured into custom molds and cooled to room temperature. Acoustic activation and data acquisition were performed using the L12-3v transducer.

Fluorescence microscopy was used to validate the distribution of ND-labeled macrophages within the agarose matrix. For this validation, 22.5×10^6^ ND-labeled macrophages in 1 mL PBS were embedded in agarose gel as described above, from which thin slices were manually sectioned and placed onto glass slides. DAPI mounting medium was applied to each slide, and the samples were imaged using the Echo Revolution microscope at 4x and 20x magnification.

### Comparison of ND activation using L12-3v and L22-14vX transducers

To assess the feasibility of using high-resolution ultrasound for *in vivo* imaging with ND-labeled macrophages, the blinking response of NDs activated by the L22-14vX transducer was compared to that induced by the L12-3v. For this comparison, NDs were embedded in two separate agarose gels as described previously, yielding final concentrations of 8.8×10^8^ NDs/mL. The phantoms were placed at the respective focal points of each transducer, and acoustic activation and data acquisition were performed to evaluate the resulting signal yield.

### Ultrasound imaging of ND-labeled MΦs in an *in vivo* tumor model

All animal procedures were performed according to the guidelines of the Institutional Animal Research Ethical Committee (approvals TAU-MD-IL-2603-120-3 and TAU-MD-IL-2605-129-5). Female BALB/c mice, aged between 8 to 12 weeks and weighing 20– 25 g (sourced from Envigo, Jerusalem, Israel), were used as the breast cancer animal model. 4T1 murine mammary carcinoma cells were cultured in Roswell Park Memorial Institute (RPMI) 1640 medium, supplemented with 10% (v/v) FBS and 1% (v/v) penicillin-streptomycin and maintained at 37 °C within a 5% CO_2_ humidified incubator. By the day of injection, they had achieved approximately 85% confluency. Cells were collected via dissociation with TrypLE and were subsequently suspended at a concentration of 9×10^5^ cells in 25 µL PBS+/+. This cell suspension was then used for subcutaneous injection into the mammary fat pad, creating the primary tumor model. Tumors were allowed to grow until they reached a size of ∼7 mm.

ND-labeled macrophages were prepared as described previously in T75 tissue culture flasks, dissociated with a cell scraper in 7 mL DMEM, quantified, centrifuged and resuspended. For ultrasound imaging of the ND-labeled macrophages *in vivo*, the mice were divided into two groups: the control group was injected with macrophages without NDs and the second with the ND-labeled macrophages. The tumors were placed in the focal point of the L22-14vX transducer, a needle was inserted intratumorally guided via ultrasound until clearly visualized as a high-intensity signal. A final load of 8×10^5^ macrophages in 40 µL was injected intratumorally. Subsequently, the acoustic activation and data acquisition were performed using the L22-14vX transducer.

### Ultrasound imaging of ND-labeled MΦs in the liver

To evaluate the biodistribution and liver accumulation of ND-labeled macrophages within an animal model, six BALB/c mice (8 to 12 weeks old, 20-25 g, Envigo, Jerusalem, Israel) were utilized.

ND-labeled macrophages were prepared as described previously in T75 tissue culture flasks, dissociated with a cell scraper in 7 mL DMEM, quantified and centrifuged. The cell pellet was resuspended to a final load of 1×10^7^ cells in 150 µL PBS for systemic administration, and a separate control group received no injection. Following a four-hour circulation period to allow for macrophage homing, the mice were sacrificed, and the livers and brains were harvested.

To detect the accumulated macrophages, ultrasound acquisition was applied to the excised organs within a custom 1.5% (w/v) agarose phantom. The phantom was cast in a mold (65 mm × 25 mm × 28 mm) containing a centrally positioned aluminum rod (24 mm height, 10 mm diameter) to create a sample well. After removal from the mold, the phantom was aligned with the focal zone of the transducer. The organs were positioned individually within the rod-shaped cavity, which was rinsed between samples. Acoustic activation and data acquisition were performed using the L22-14vX transducer.

## Acknowledgments

This work was supported in part by the Israel Science Foundation under Grant No. 192/22, in part by an ERC StG under Grant No.101041118 (NanoBubbleBrain), the Israel Cancer Research Fund (GrantNo. 1286686), and in part by the Nicholas and Elizabeth Slezak Super Center for Cardiac Research and Biomedical Engineering at Tel Aviv University. S. Gotshal Zahavi would like to acknowledge her funding via the Marian Gertner Institute for Medical Nanosystems.

## Author contribution statement

S. Gotshal Zahavi designed and performed the research, analyzed the data, and wrote the paper. M. Bismuth and T. Bercovici assisted in experiments. T. Ilovitsh guided, advised, and designed the research and wrote the paper.

## Competing interest statement

The authors declare no conflict of interest.

## Data availability

The datasets generated during and/or analyzed during the current study will be made available upon request.

## Notes

### Competing Interest Statement

The authors have declared no competing interest.

